# Untangling cortical maps in mouse primary visual cortex

**DOI:** 10.1101/102079

**Authors:** Luis O. Jimenez, Elaine Tring, Joshua T. Trachtenberg, Dario L. Ringach

## Abstract

Local populations of neurons in mouse visual cortex exhibit diverse tuning preferences. We show this seeming disorder can be untangled — the similarity of tuning between pairs of neurons is correlated better with the overlap between their receptive fields in visual space rather than with their distance in the cortex. These findings are consistent with the hypothesis that salt-and-pepper maps arise from the lateral dispersion of clonally related neurons.

---

In mouse primary visual cortex (V1) clonally related cells derived from a single progenitor tend to have similar tuning properties and are more likely to connect with each other. This observation suggests that cell lineage is involved in the development of cortical microcircuits^1,2^. Furthermore, it has been proposed that salt-and-pepper maps in mice may arise due to lateral dispersion of sister neurons in the cortex, while a tighter distribution along radial columns in primates could generate orderly maps^3^.

To appreciate the implications of this idea consider a model where colonally related cells (or “sister neurons”), known to be transiently coupled via gap junctions during the first postnatal week^4^, come to share common feedforward inputs that endows them with similar tuning preferences^5^. As the location of a neuron’s receptive field in visual space is primarily determined by its feedforward inputs, one would expect sister neurons to develop overlapping receptive fields as well. As we show below, this prediction holds true.

Let us further assume, for the sake of discussion, that the thalamic input to the cortex is organized retinotopically and that local biases in the input imprint a feature preference map early in development (Fig 1). Such biases may originate from a limited number of local ON/OFF inputs^6-8^ or from individual thalamic afferents that are tuned^9^. Two cortical organizations are possible depending on the adult distribution of sister cells within the cortex. In one scenario, sister neurons could arrange themselves along radial columns, leading to a columnar architecture and maps of stimulus preference and retinotopy (Fig 1a). Here, any relationship between tuning and retinotopy present at the input, *θ* = *f*(*x_v_*), is very well preserved in the horizontal dimension of the cortex (up to a magnification factor), *θ* ≅ *f*(*x_c_*), due to the precise radial arrangement of sister neurons (Fig 1a).

**Figure 1.**
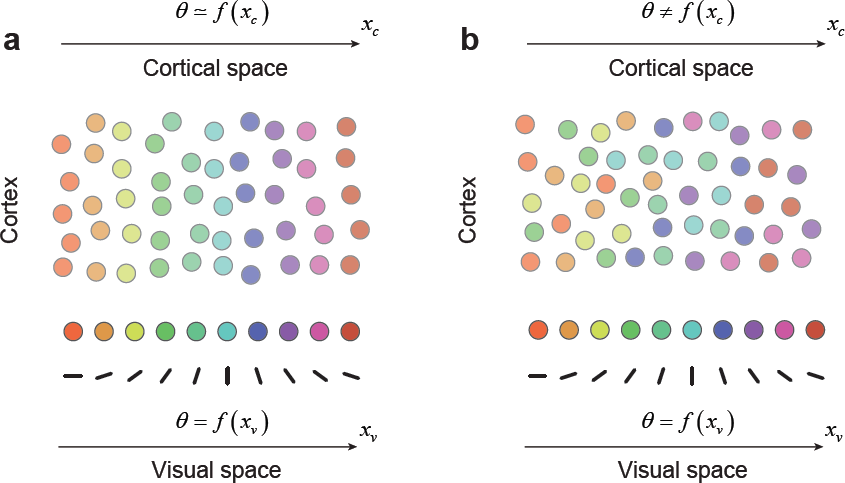
Cell lineage and dispersion of clonally related neurons as a major determinant of cortical organization. (**a**) A scenario where clonally related neurons are distributed radially in the cortex with little or no lateral dispersion, resulting in orderly retinotopic and feature maps. (**b**) A scenario where clonally related neurons distribute radially and laterally in the cortex, resulting in a salt-and-pepper map and a high local scatter of receptive field positions.

Alternatively, sister neurons could disperse laterally in the cortex and intermingle with neurons originating from other progenitor cells (Fig 1b). This would result is a salt-and-pepper map of tuning preferences and a concomitant increase in the scatter of their receptive field positions, which are key features of mouse V1^6,10^. Strong spatial mixing caused by lateral dispersion would significantly weaken or obliterate any relationship between tuning and retinotopy. However, the latent organization of the input, *θ* = *f*(*x_v_*), could still be recovered by measuring the tuning and receptive field location of neurons in the population, as this relationship remains unaffected by the spatial mixing (Fig 1b). Here we show that it is possible to recover hidden spatial structure in mouse V1 by “untangling” the putative effects of lateral dispersion.

With this simple model as background we studied the joint organization of tuning and retinotopy in primary visual cortex by means of resonant, two-photon microcopy in alert mice expressing GCaMP6f in the superficial layers of V1 (Fig 2a). To measure the tuning of neurons to orientation and spatial frequency (the Fourier domain) we used a visual stimulus consisting of a long sequence of flashed, high-contrast sinusoidal gratings that had random orientations and spatial frequencies (Fig 2b, top). We estimated the tuning of each cell in the joint spatial frequency and orientation domain by linearly regressing the response on the stimulus^11^ (Fig 2c). We denote the resulting tuning kernel of the *i* − *th* cell in the population by 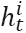. Similarly, we measured the location of receptive fields in the visual field by using a sparse noise stimulus in which dark and bright spots (5 degrees diameter) were randomly flashed on the computer screen for a brief period of time before being relocated (Fig 2b, bottom). We computed separate ON and OFF maps for each cell by reverse correlating their responses with the location of the bright and dark disks respectively (Fig 2d)^12^. We denote the maps corresponding to the *i* − *th* cell in the population by 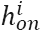 and 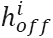.

**Figure 2.**
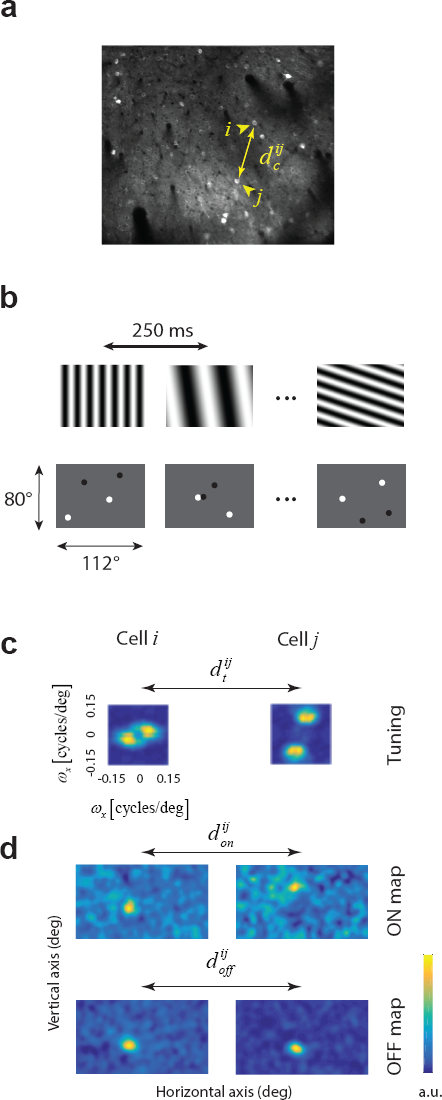
Measuring tuning and ON/OFF sub-region locations in mouse primary visual cortex. (**a**) GCaMP6f was expressed in excitatory neurons of layer 2+3 in primary visual cortex. The distance between neural pairs was defined as their distance in the imaging plane. (**b**) Pseudo-random sequences of full-field, sinusoidal gratings were used to map the tuning for orientation and spatial frequency. Sparse noise stimuli consisting of randomly flashed dark and bright disks was used to map the ON and OFF sub-regions of each cell. (**c**) Two examples of tuning kernels in the Fourier domain (top). Here, the origin is at the center of the image. The distance of a given point on the image to the origin represents its spatial frequency, and its angle with respect to the positive *x* axis represents the orientation^11,17^. The kernels are normalized and presented in arbitrary units. Maximal spatial frequency along the *x* and *y* axes was 0.15 cycles/deg. (**d**) Two examples of tuning kernels in the Fourier domain (top). Here, the origin is at the center of the image. The distance of a given point on the image to the origin represents its spatial frequency, and its angle with respect to the positive *x* axis represents the orientation. The kernels are normalized and presented in arbitrary units. Maximal spatial frequency along the *x* and *y* axes was 0.15 cpd. (**e**) Two examples of ON/OFF kernels. The dimensions of the image in visual space correspond to the ones displayed in (**b**).

We define the tuning similarity between cell *i* and cell *j* as the correlation coefficient between their corresponding tuning kernels, 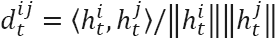 (Fig 2c). Similarly, we define the overlap between ON/OFF sub-regions, 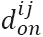 and 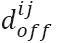, as the respective correlation coefficients of their maps (Fig 2c).

Some cells had only a significant ON map, a significant OFF map, or both. We define the subset of neuron pairs where we could compute both 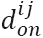 and 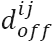 as *S*_*on*+*off*_. Subsets of neuron pairs where only ON or OFF overlap could be measured are denoted by *S_on_* and *S_off_* respectively. Only cells with good tuning kernels were selected for analysis, meaning that we could compute 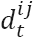 in all cases. The physical distance between two cells in imaging plane is denoted by 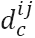 (Fig 2a). Cell pairs were pooled over 19 non-overlapping imaging fields in 11 mice.

Consistent with local diversity^13^ we find no significant correlation between tuning similarity and cortical distance (Fig 3a, Table1) (but see ref^11^). Interestingly, as predicted by the lateral dispersion model, there is a relationship between tuning selectivity and receptive field overlap (Fig 3b, Table 1). Cells with high overlap tend to have high tuning similarity; this is particularly the case for cell pairs that share ON and OFF sub-regions (Fig 3b, solid symbols). In contrast, pairs of cells with non-overlapping subregions can attain a range of tuning similarities values, some of which can be high, but not as high as cells that share ON and OFF sub-regions. Notably, there is a scarcity of cell pairs which have high overlap but disparate tuning (Fig 2b), and that the overlap between ON and OFF sub-regions correlates (Fig 3c). Both of these features are consistent with the presence of local bias or constraints in the thalamic inputs. Finally, a more orderly map of preferences reveals itself when plotting the tuning kernels as a function of their receptive field locations in the visual field (Fig 3e) rather than in cortical space (Fig 3d). The highlighted sets {*a, c*}, {*b, d, j, e, h*}, and {*f, g*}, share similar orientation preferences and cluster in visual space, but are intermingled in cortical space. This outcome is a consequence of the correlation between similarity of tuning and receptive field overlap (Fig 3b). We are in the process of collecting larger datasets from individual animals to perform a denser reconstruction of such putative maps in mouse V1 and gain further insights into their two-dimensional organization.

**Figure 3.**
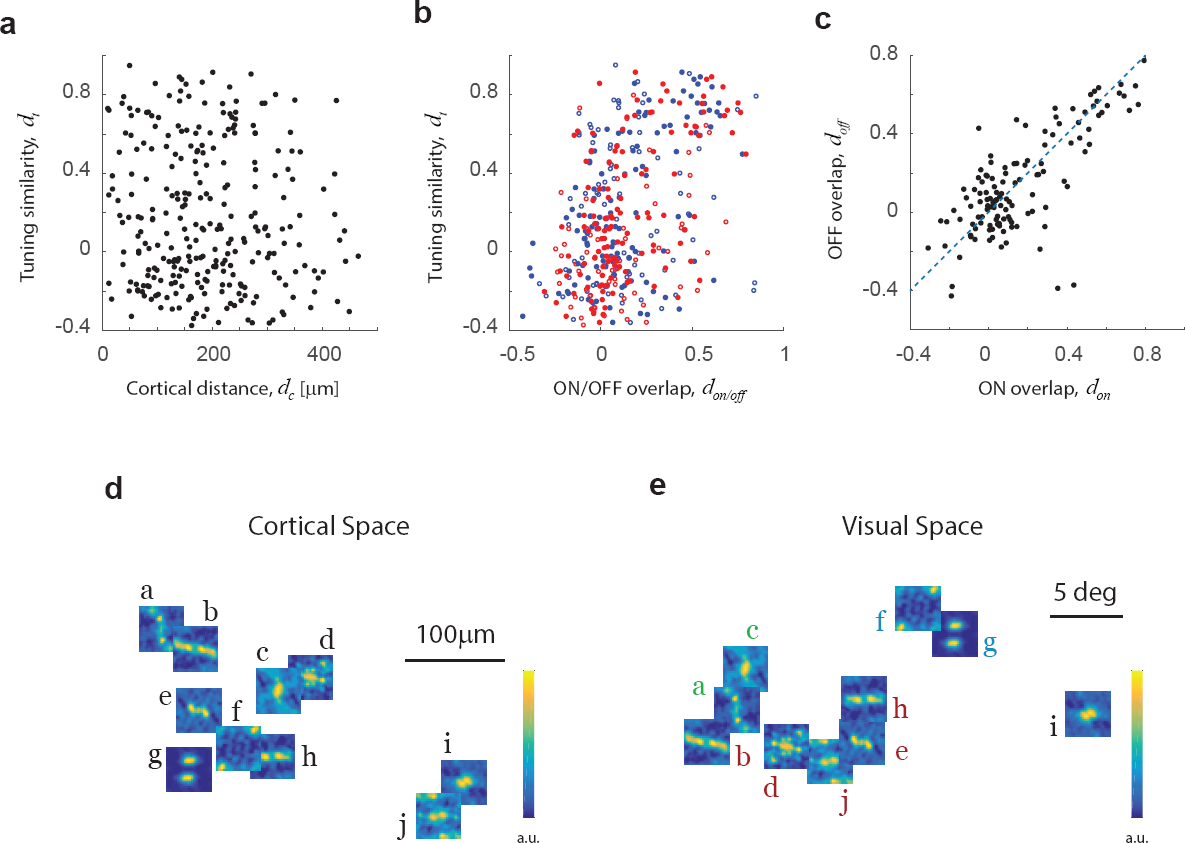
Untangling cortical maps. (**a**) There is no statistical correlation between similarity of tuning and cortical distance. (**b**) There is a strong statistical correlation between similarity of tuning and overlap of receptive fields in visual space. Solid dots indicate cell pairs which had both ON and OFF maps. Red dots show overlap of ON sub-regions; blue dots show overlap of OFF sub-regions. Open symbols show pairs where only ON (red) or OFF (blue) sub-region overlap could be computed (see Table 1 for statistical tests). (**c**) For cell pairs where both ON and OFF maps could be computed there is a strong correlation between their degree of overlap (see Table 1 for statistical tests). (**d**) Kernels displayed as a function of the location of cell bodies in the cortex shows a highly variable organization. (**e**) When the same kernels are plotted as a function of their location in visual space, kernels with similar tuning cluster together.

**Table 1.** Statistical models were used to predict tuning similarity based on cortical distance, overlap of ON and OFF sub-regions. There was no significant correlation between tuning similarity and cortical distance, but a highly significant effect with both ON and OFF sub-region overlap. Moreover, the overlap of ON and OFF sub-regions were highly and significantly correlated.

| Domain | Model | Coefficient | Significance | *n* | Fig |
| --- | --- | --- | --- | --- | --- |
| *All cell pairs* | $d_t \sim \alpha + \beta d_c$ | $\beta = -0.002$ | $p = 0.35$ | 294 | **3a** |
| $S_{on} \cup S_{on+off}$ | $d_t \sim \alpha + \beta d_{on}$ | $\beta = 0.71$ | $p = 4 \times 10^{-13}$ | 190 | **3b** |
| $S_{off} \cup S_{on+off}$ | $d_t \sim \alpha + \beta d_{off}$ | $\beta = 0.09$ | $p = 6 \times 10^{-9}$ | 210 | **3b** |
| $S_{on+off}$ | $d_t \sim \alpha + \beta d_{on} + \gamma d_{off}$ | $\beta = 0.47, \gamma = 0.44$ | $p_\alpha = 0.004, p_\beta = 0.005$ | 125 | **3b** |
| $S_{on+off}$ | $d_{off} \sim \alpha + \beta d_{on}$ | $\beta = 0.77$ | $p = 7.8 \times 10^{-24}$ | 125 | **3c** |

The present findings are consistent with idea that are orderly, latent maps of feature selectivity and retinotopy are present early in development^3^. In the mouse, this spatial organization appears disrupted by the lateral dispersion of sister cells, but may leave some traces of organization behind^11,14^. Here we showed that the underlying organization, if it exists, ought to reveal itself when analyzing the relationship between preference in visual space, instead of cortical space (Fig 3d, e). Finally, we note the lateral dispersion model (Fig 1), de-emphasizes the importance of cortical self-organization mechanisms in guiding map formation^15^, while favoring hypotheses that put forward a link between tuning and visual space early in development^7,16^.

## Methods

### Animals

All procedures were approved by UCLA’s Office of Animal Research Oversight (the Institutional Animal Care and Use Committee), and were in accord with guidelines set by the US National Institutes of Health. A total of 11 C57BL/6J mice (Jackson Laboratory), both male (5) and female (6), aged P35-56, were used in this study. Mice were housed in groups of 2-3, in reversed light cycle. Animals were naïve subjects with no prior history of participation in research studies. We imaged 19 different fields to obtain the data discussed in this paper.

### Surgery

Carprofen and buprenorphine analgesia were administered pre-operatively. Mice were then anesthetized with isoflurane (4-5% induction; 1.5-2% surgery). Core body temperature was maintained at 37.5C using a feedback heating system. Eyes were coated with a thin layer of ophthalmic ointment to prevent desiccation. Anesthetized mice were mounted in a stereotaxic apparatus. Blunt ear bars were placed in the external auditory meatus to immobilize the head. A portion of the scalp overlying the two hemispheres of the cortex (approximately 8mm by 6mm) was then removed to expose the underlying skull.

After the skull is exposed it was dried and covered by a thin layer of Vetbond. After the Vetbond dries (approximately 15 min) it provides a stable and solid surface to affix the aluminum bracket with dental acrylic. The bracket is then affixed to the skull and the margins sealed with Vetbond and dental acrylic to prevent infections.

### Virus injection

A 3mm diameter region of skull overlying the occipital cortex was removed. Care was taken to leave the dura intact. GCaMP6-fast (UPenn Vector Core: AAV1.Syn.GCaMP6f.WPRE.SV40; #AV-1-PV2822) was expressed in cortical neurons using adeno-associated virus (AAV). AAV-GCaMP6-fast (titer: ~10ˆ13 genomes ml^-1^) was loaded into a glass micropipette and slowly inserted into the primary visual cortex (V1) using a micromanipulator. Two injection sites were made centered around the center of V1 and separated about 200 microns apart. For each site, AAV-GCaMP6-fast was pressure injected using a PicoSpritzer III (Parker, Hollis, NH) (4 puffs at 15-20 pounds per square inch with a duration of 10 ms, each puff was separated by 4 s) starting at a depth of 350 microns below the pial surface and making injections every 10 microns moving up with the last injection made at 100 microns below the pial surface. The total volume injected across all depths was approximately 0.5 μl. The injections were made by a computer program in control of the micro-manipulator and the Picosprtizer.

A sterile 3mm diameter cover glass was then placed directly on the dura and sealed at its edges with VetBond. When dry, the edges of the cover glass were further sealed with dental acrylic. At the end of the surgery, all exposed skull and wound margins were sealed with VetBond and dental acrylic and a small, sealed glass window was left in place over the occipital cortex. Mice were then removed from the stereotaxic apparatus, given a subcutaneous bolus of warm sterile saline, and allowed to recover on the heating pad. When fully alert they were placed back in their home cage.

### Imaging

Once expression of Gcamp6f was observed in primary visual cortex, typically between 11-15 days after the injection, imaging sessions took place. Imaging was performed using a resonant, two-photon microscope (Neurolabware, Los Angeles, CA) controlled by Scanbox acquisition software (Scanbox, Los Angeles, CA). The light source was a Coherent Chameleon Ultra II laser (Coherent Inc, Santa Clara, CA) running at 920nm. The objective was an x16 water immersion lens (Nikon, 0.8NA, 3mm working distance). The microscope frame rate was 15.6Hz (512 lines with a resonant mirror at 8kHz). Eye movements and pupil size were recorded via a Dalsa Genie M1280 camera (Teledyne Dalsa, Ontario, Canada) fitted with a 740nm long-pass filter that looked at the eye indirectly through the reflection of an infrared-reflecting glass (Fig 1a). Images were captured at an average depth of 200 μm.

### Visual stimulation

Hartley^18,19^ and sparse noise stimuli were generated in real-time by a Processing sketch using OpenGL shaders (see http://processing.org). The Hartley stimulus was updated 4 times a second on a BenQ XL2720Z screen refreshed at 60Hz. The screen measured 60 cm by 34 cm and was viewed at 20 cm distance, subtending 112 × 80 degrees of visual angle. The maximum spatial frequency was 0.15 cycles/deg. Sparse noise consisted of flashed dark and bright disks with a diameter of 5 deg. Two disks of each contrast appeared at any one time. The lifetime of each disk was 250 ms, after which it was removed and repositioned randomly on the screen. TTL signals generated by the stimulus computer were sampled by the microscope and time-stamped with the frame and line number that being scanned at that time. The time-stamps provided the synchronization between visual stimulation and imaging data.

The screen was calibrated using a Photo-Research (Chatsworth, CA) PR-650 spectro-radiometer, and the result used to generate the appropriate gamma corrections for the red, green and blue components via an nVidia Quadro K4000 graphics card. The contrast of the stimulus was 80%. The center of the monitor was positioned with the center of the receptive field population for the eye contralateral to the cortical hemisphere under consideration. The location of the receptive fields were estimated by an automated process where localized, flickering checkerboards patches, appeared at randomized locations within the screen. This experiment was ran at the beginning of each imaging session to ensure the centering of receptive fields on the monitor.

### Image processing

The image processing pipeline was the same as described in detail elsewhere11. Briefly, calcium images were aligned to correct for motion artifacts. Following motion stabilization, we used a Matlab graphical user interface (GUI) tool developed in our laboratory to manually define regions of interest corresponding to putative cell bodies. Following segmentation, we extracted signals by computing the mean of the calcium fluorescence within each region of interest and discounting the signals from the nearby neuropil.

### Kernel estimation

The estimation of the tuning kernel was performed as in earlier studies by fitting a linear model between the response and the stimulus11. Kernels that had no structure often showed a distribution of kernel coefficients that was close to normal. Thus, we imposed a minimum kurtosis value of 4, which selected kernels that had some discernible structure. Similarly, we computed the ON/OFF maps by fitting a linear model between the response and the location of bright and dark spots. The maps were smoothed with the Gaussian window of σ = 5 deg. We declared spatial maps with coefficient distributions larger than 4 as showing significant spatial structure, otherwise we declared the cell unresponsive to the corresponding contrast. The outcome of the present analyses was robust to the selection of the threshold for kurtosis.

## Code and data availability

All analyses were conducted in Matlab (Mathworks, Natick, MA). The code and data are available upon request from the authors.

## Contributions

ET performed surgeries and ran experiments. LOJ performed analyses. JTT consulted on developmental models and wrote manuscript. DLR designed and performed the experiment, executed the analyses, wrote the manuscript, and supervised the project.

